# The fungal root endophyte *Serendipita indica* modifies extracellular nucleotides to subvert plant immunity

**DOI:** 10.1101/396028

**Authors:** Shadab Nizam, Xiaoyu Qiang, Stephan Wawra, Robin Nostadt, Felix Getzke, Florian Schwanke, Ingo Dreyer, Gregor Langen, Alga Zuccaro

## Abstract

**One sentence abstract:** Immune modulation by metabolites in plant fungus interaction

**Abstract:** Extracellular adenosine 5′-triphosphate (eATP) is an essential signaling molecule that mediates different cellular processes through its interaction with membrane-associated receptor proteins in animals and plants. eATP regulates plant growth, development and responses to biotic and abiotic stresses. Its accumulation in the apoplast induces ROS production and cytoplasmic calcium increase mediating a defense response to invading microbes. We demonstrate that perception of eATP is important in plant-fungus interaction and that during colonization by the beneficial root endophyte *Serendipita indica* accumulation of eATP in the apoplast occurs at early symbiotic stages. We show by liquid chromatography-tandem mass spectrometry, cytological and functional analysis that *S. indica* subvert eATP host response by secreting *Si*E5’NT, an enzymatically active ecto-5′nucleotidase capable of hydrolyzing eATP to adenosine. *A. thaliana* lines producing extracellular *Si*E5’NT are signi?cantly better colonized and have reduced eATP levels and defense signaling, indicating that *Si*E5’NT functions as a compatibility factor. Our data show that extracellular bioactive nucleotides play an important role in fungus-root interactions and that fungi can modify plant derived metabolites in the apoplast to modulate host immunity.

## Introduction

ATP is a coenzyme that serves as an universal energy currency and as building block of genetic material and secondary metabolites. Additionally, ATP and adenosine are important extracellular regulators in plants and animals (Burnstock, 1972, Burnstock, 2012, Chivasa & Slabas, 2012, Clark, Morgan et al., 2014, Tanaka, Choi et al., 2014). In plants it has been shown that extracellular ATP (eATP) is released during growth, physical injury and in response to various abiotic and biotic stresses and microbe derived elicitors (Cao, Halane et al., 2017, Choi, Tanaka et al., 2014, Clark et al., 2014, Dark, Demidchik et al., 2011, Kim, Sivaguru et al., 2006, Roux & Steinebrunner, 2007, Song, Steinebrunner et al., 2006, Tanaka et al., 2014, Tanaka, Gilroy et al., 2010, Wu & Wu, 2008). Accumulation of eATP triggers the production of reactive oxygen species (ROS), nitric oxide, callose deposition, cytoplasmic calcium increase, transient phosphorylation of MPK3 and MPK6, and expression of genes involved in plant stress response and immunity (Choi et al., 2014, Lim, Wu et al., 2014). To be physiologically relevant eATP must be sensed by specific receptors at the cell surface. Recently, the *Arabidopsis thaliana* lectin receptor kinase DORN1 (DOes not Respond to Nucleotides 1, At5g60300), also known as the LecRK-I.9 (lectin receptor kinase I.9), was identified as the first plant purinoceptor essential for the plant response to eATP, unveiling an eATP signaling pathway in plants (Choi et al., 2014). The RXLR-dEER effector protein IPI-O secreted by the oomycete pathogen *Phytophthora infestans* targets LecRK-I.9/DORN1 (Bouwmeester, de Sain et al., 2011, Gouget, Senchou et al., 2006, Senchou, Weide et al., 2004, Wang, Nsibo et al., 2016), suggesting that some microbes have the tools to manipulate host eATP perception/signaling. LecRK-I.9/DORN1 mutant plants show enhanced susceptibility to pathogenic infection by the oomycete *P. brassicae* and the bacterium *Pseudomonas syringae* (*Pst*). Consistently, overexpression of LecRK-I.9/DORN1 increases plant resistance to *Phytophthora* spp and *Pst* (Balague, Gouget et al., 2017). Taken together, these results suggest a role for an eATP-receptor protein in plant immunity.

There is increasing evidence that fungi, both pathogenic and mutualistic, have large repertoires of secreted effectors. Some were functionally shown to be able to manipulate the host cell physiology, suppress plant defense and ultimately promote fungal colonization (Rovenich, Boshoven et al., 2014). Among fungi, compatibility factors are mainly reported from biotrophic and hemibiotrophic leaf pathogens (Lo Presti, Lanver et al., 2015). In contrast only few factors have been functionally characterized from root mutualistic fungi (Kloppholz, Kuhn et al., 2011, Plett, Daguerre et al., 2014, Plett, Kemppainen et al., 2011, Wawra, Fesel et al., 2016). Therefore, the basis and mode of action of mutualistic fungal compatibility factors remain poorly understood. The filamentous root endophyte *Serendipita indica* (syn. *Piriformospora indica*) belongs to the order *Sebacinales* (Basidiomycota). *S. indica* colonizes the root epidermal and cortex cells without penetrating the central cylinder and displays a biphasic colonization strategy (Deshmukh, Huckelhoven et al., 2006, Lahrmann, Ding et al., 2013, Qiang, Zechmann et al., 2012b, Zuccaro, Lahrmann et al., 2011). During the initial phase of biotrophic colonization, the fungus invades the root cells inter-and intracellularly. Subsequently, *S. indica* switches to a host cell death associated phase, although a defined switch to necrotrophy with massive cell death does not occur (Deshmukh et al., 2006, Lahrmann et al., 2013, Qiang et al., 2012b). *S. indica* colonization exhibits various effects on the host plants such as enhanced growth and assimilation of nitrate and phosphate, increased tolerance to abiotic stresses and resistance against pathogens (Lahrmann & Zuccaro, 2012, Oberwinkler, Riess et al., 2013, Qiang, Weiss et al., 2012a, Vahabi, Sherameti et al., 2015). Since *S. indica* establishes symbiotic interactions with a wide range of experimental plants, including the dicot model plant *A. thaliana* and the monocot cereal crop *Hordeum vulgare* (barley), it represents an excellent model system to study ATP-mediated plant defense responses in the roots of unrelated hosts.

In order to identify *S. indica* secreted effectors, proteins present in the apoplastic fluid (APF) of colonized barley roots were analyzed at three different symbiotic stages by liquid chromatography-tandem mass spectrometry (LC-MS/MS). One of the most abundant secreted fungal proteins identified at all time points is a predicted 5’-nucleotidase. The gene encoding this enzyme is induced during colonization of both, barley and *A. thaliana*. Animal ecto-5’-nucleotidases (E5’NTs) have been considered to play a key role in the conversion of AMP to adenosine and in counteracting together with ectonucleotide pyrophosphatase/phosphodiesterase (E-NPP), ecto-nucleoside triphosphate diphosphohydrolase (E-NTPDase) and alkaline phosphatases (AP), eATP signaling (Antonioli, Pacher et al., 2013, Matsuoka & Ohkubo, 2004). The importance of bioactive nucleotides signaling and fungal extracellular E5’NT activity during plant-fungus interaction is unknown. We show here that *S. indica* E5’NT is capable of hydrolyzing ATP, ADP and AMP to adenosine, altering eATP levels and defense signaling *in planta* and plays an important role in fungal accommodation at early symbiotic stages, suggesting that immune modulation by metabolites in the apoplast is a mechanism of compatibility in plant-fungus interactions.

## Results

### Identification of fungal proteins in the apoplast of barley roots

In order to identify soluble secreted candidate effector proteins during *S. indica* root colonization, the proteins present in the APF of barley roots at three different symbiotic stages (biotrophic and early and late cell death associated phase corresponding to 5, 10 and 14 day-post-inoculation respectively) were analyzed together with the proteins found in the culture filtrate (CF) obtained from *S. indica* axenically grown in liquid complex medium (CM). To assess possible cytoplasmic contaminations a strain constitutively expressing an *S. indica* codon-optimized GFP under the *Si*GPD promoter (Hilbert, Voll et al., 2012b) was used. Absence of fungal cytoplasmic GFP protein contamination due to cell lysis was corroborated by western blotting of the apoplastic protein samples using anti-GFP antibody (Fig. S1A).

LC-MS/MS analysis after tryptic digestion led to the identification of 102 *S. indica* proteins putatively targeted to the apoplast of which thirty-three were present at all three time points in at least one of the biological replicates (Table S1 and Fig. 1A, B). Predictions using ApoplastP (http://apoplastp.csiro.au/) and SecretomeP (http://www.cbs.dtu.dk/services/SecretomeP-1.0/) indicated that 48 of the 102 *S. indica* proteins are putatively targeted to the plant apoplast (47%) with 9 proteins predicted to be secreted via a non canonical secretion pathway using a cutoff of 0.6 (Table S2). No peptides for GFP were found in any of the apoplastic fluid samples, confirming the GFP western blotting analysis. Twenty proteins were unique at 5 dpi, 4 at 10 dpi and 21 at 14 dpi, suggesting differential secretion of *S. indica* proteins at different stages of colonization (Table S1).

**Figure 1.**
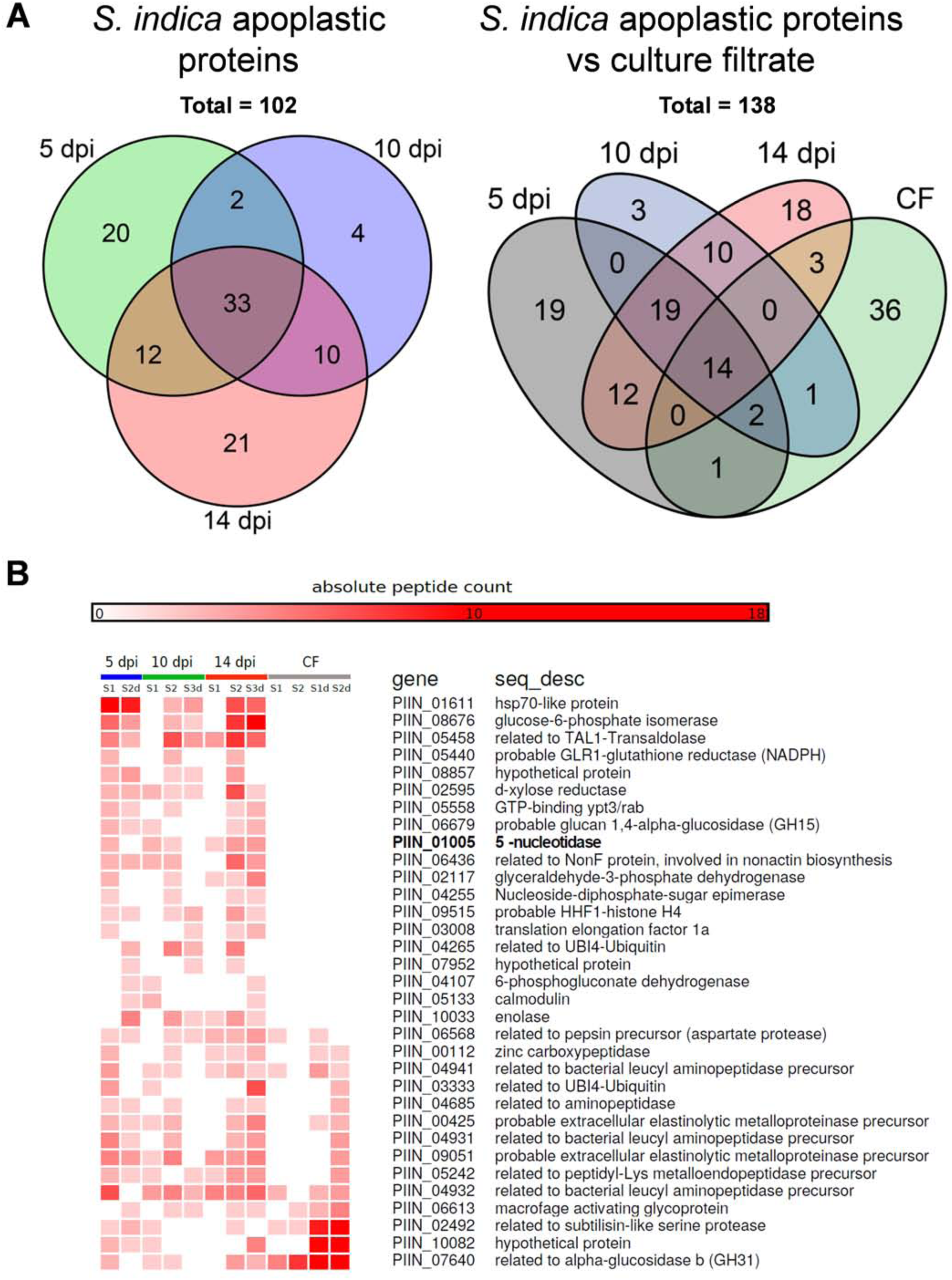
Identification of *S. indica* apoplastic proteins. **A** Distribution of *S. indica* apoplastic proteins identified by LC-MS/MS analysis from different symbiotic stages (5, 10 and 14 dpi) and in relation to proteins identified in CF of *S. indica* axenically grown in CM. In total, 102 *S. indica* proteins were identified in the APF of colonized barley roots. Of these, 33 proteins were present at all three time points. Twenty proteins were unique at 5 days post infection (dpi), 4 at 10 dpi and 21 at 14 dpi. **B** Heat map showing absolute counts of unique peptides for *S. indica* apoplastic proteins present at all-three time points and in culture filtrate.

Twenty one of the 102 identified apoplastic proteins were also found in the culture filtrate of *S. indica* grown in CM, which included a large proportion of enzymes acting on peptide bonds (Fig. 1A, B). The majority of these proteases might conceivably play a role in nutrition rather than suppression of host defense because of the large overlap with the culture filtrate proteome and because amino acid uptake seem to be an active process during root colonization based on transcriptomic data (Lahrmann et al., 2013, Lahrmann, Strehmel et al., 2015, Zuccaro et al., 2011). Four peptidases (PIIN_00628, PIIN_02239, PIIN_02952, PIIN_07002) however were specifically present in the apoplastic fluid which make them good candidate effectors possibly involved in host defense suppression (Table S1).

Remarkably, 19 of the 33 common proteins found in the apoplastic fluid at all 3 time points where not present in the CF samples, implying an induced secretion for these proteins during interaction with the plant roots (Table S1, and Fig. 1B). Recently it was reported that *Rhizobium radiobacter* (syn. *Agrobacterium tumefaciens*, syn. *Agrobacterium fabrum*) is an endofungal bacterium of *S. indica* and that together form a tripartite Sebacinalean symbiosis with a broad range of plants (Guo, Glaeser et al., 2017, Sharma, Schmid et al., 2008). No peptides for bacterial-derived proteins were found in the apoplastic fluid of colonized barley roots and in the CF, suggesting that *R. radiobacter*´s contribution to successful symbiosis does not involve accumulation of bacterial proteins in the apoplast of barley.

In a first step for functional interpretation of the resultant protein list from the apoplastic fluid we assigned Gene Ontology terms to the identified proteins (Conesa, Gotz et al., 2005, Supek, Bosnjak et al., 2011, Zheng & Wang, 2008). GO-term enrichment analysis for biological processes showed that in the culture filtrate obtained from axenically grown *S. indica* there is an over representation of GO-terms involved in proteolysis whereas the GO enriched terms of apoplastic fungal proteins were related to purine metabolism, and specifically to ATP metabolic processes (Table S3, Fig. S1B and S2).

Among the fungal proteins specifically present in the apoplastic fluid of colonized roots, several ATP-scavenging enzymes, including a 5’-nucleotidase (PIIN_01005) and a nucleoside diphosphate kinase (PIIN_00784) were identified (Table S1 and Fig. S1B, C). Twenty one orthologues of the *S. indica* apoplastic proteins were also found in the apoplastic fluid of rice leaves infected by the rice-blast pathogen *Magnaporthe oryzae* (Table S4 and Fig. S1B asterisks) (Kim, Wang et al., 2013). Four of these proteins are predicted enzymes involved in ATP metabolism. The presence of fungal derived ATP-scavenging enzymes in the apoplast of colonized plants indicates that extracellular purine metabolism is possibly important in plant-fungus interaction.

### eATP plays a role during fungal colonization in barley and in Arabidopsis

The enrichment for GO terms related to ATP metabolic processes among the fungal-derived apoplastic proteins led us to test whether root colonization by *S. indica* triggers changes in eATP levels at different symbiotic stages and in different plant species. Therefore, a specific luminescent ATP detection assay was used to assess ATP levels of Arabidopsis seedlings grown in liquid medium or barley apoplastic fluid samples both from *S. indica* colonized and mock treated plants at 3, 5, 7 and 10 dpi. The results show that eATP levels are significantly higher in *S. indica* colonized roots compared to mock treated plants, especially at early symbiotic stages where the biotrophic interaction is still predominant (Fig. 2A, B). Thus, there is a bulk release/accumulation of eATP during *S. indica* biotrophic colonization of both, *Arabidopsis* and barley.

**Figure 2.**
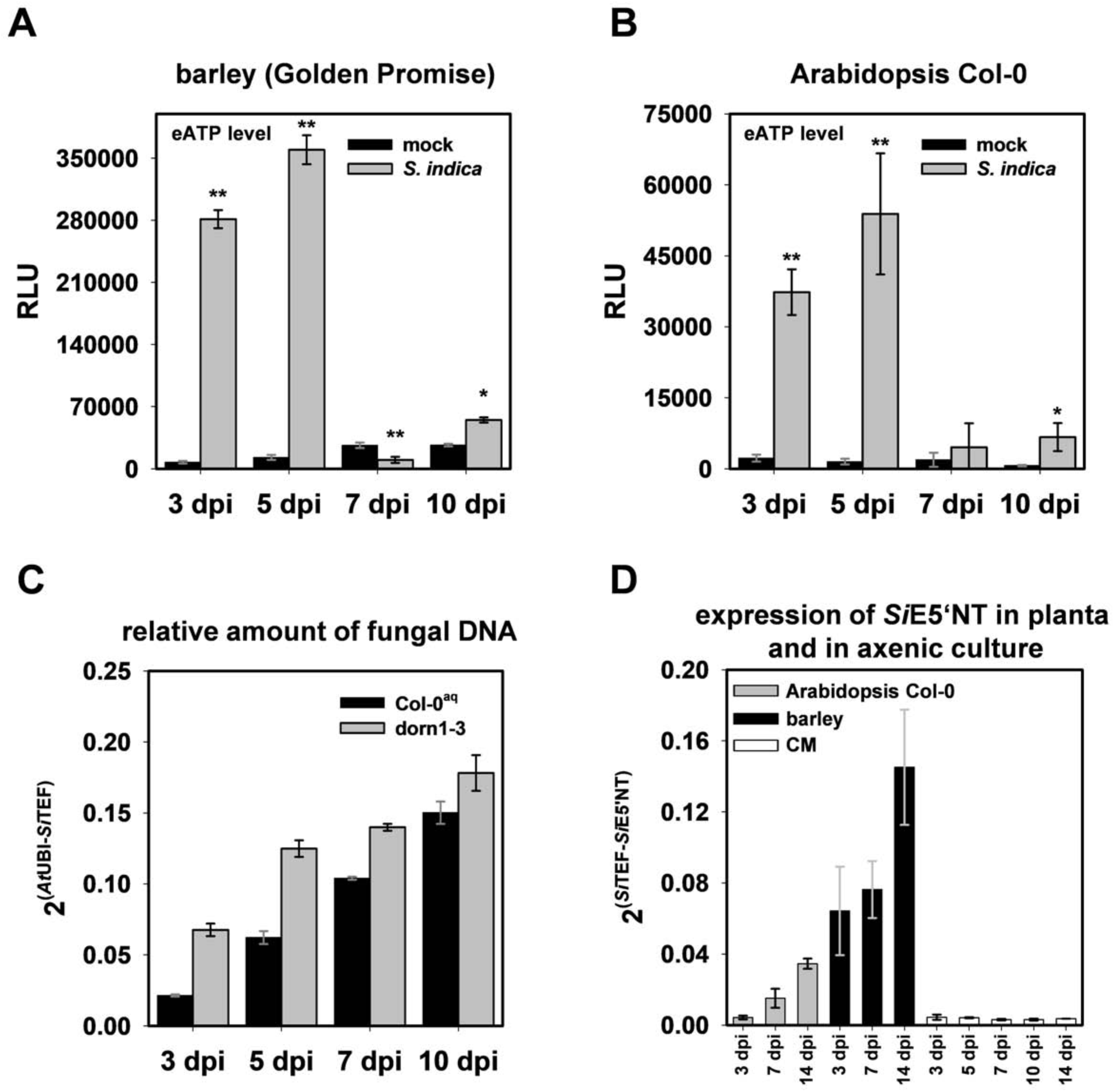
*S. indica* colonization leads to increased eATP levels in barley and *Arabidopsis* roots. **A** eATP levels in APF from *S. indica* inoculated and mock-treated barley root samples collected at the predominantly biotrophic phase (3 and 5 dpi) as well as the predominantly cell death associated phase (7 and 10 dpi). Error bars show the standard error of the mean obtained from three independent biological replicates. **B** eATP levels measured from culture medium collected from mock treated and *S. indica* colonized *Arabidopsis* seedlings at 3, 5, 7 and 10 dpi. Error bars in A and B represent ±SE of the mean from three independent biological replicates. RLU: relative light units. Asterisks indicate significance at p< 0.05 (*), 0.01 (**) analyzed by Student’s t-test. **C** *S. indica* colonization of *Arabidopsis dorn1-3* mutant and the parental Col-0^aq^ lines quantified by qPCR at 3, 5, 7 and 10 dpi. The ratio of fungal (*Si*TEF) to plant (*At*UBI) amplicons representing fungal colonization levels in plant root tissue was calculated using gDNA as template and the 2^−ΔCT^ method. Error bars represent standard error of the mean of three technical replicates. The experiment was repeated 3 times for 3, 5 and 7 dpi with similar outcomes. **D** Transcript levels of *S. indica* E5’NT during colonization of barley and Arabidopsis at different symbiotic stages and in axenic culture. Error bars represent standard error of the mean of three independent biological replicates. CM = complex medium.

Previous reports have indicated that eATP modulates plant immunity by triggering the production of ROS, calcium influx, callose deposition and expression of plant immune responsive genes (Choi et al., 2014, Lim et al., 2014). However, the role of eATP signaling during plant root-fungus interaction is still unclear. To investigate whether eATP detection plays a role during root colonization, we used an eATP-insensitive *A. thaliana* mutant, *dorn1-3* defective in the LecRK-I.9 (Choi et al., 2014). After confirming the inability of the *dorn1-3* mutant to respond to eATP and eADP with cytoplasmic calcium influx (Fig. S3), colonization of this line along with the appropriate Col-0 control (expressing the calcium reporter protein aequorin, Col-0^aq^) was assessed by qPCR at different time points (3, 5, 7, and 10 dpi). We observed that the *dorn1-3* mutant supports significant higher fungal colonization in comparison to Col-0^aq^ control, especially at early symbiotic stages (Fig. 2C and 4E). Importantly, these observations strongly indicate that eATP accumulates in the apoplast and its detection by the host affects fungal root colonization.

### S. indica PIIN_01005 is an ecto-5’-nucleotidase with broad substrate activity against ATP, ADP and AMP

Since our data show that detection of eATP plays a role during root-fungus interaction we hypnotized that *S. indica* uses some intrinsic mechanism to manipulate eATP levels to counteract the ATP-mediated host immune responses. The secreted *S. indica* PIIN_01005, consistently found in the apoplastic fluid but not in the culture filtrate samples, is a suitable candidate for such a mechanism because of its putative hydrolytic activity on purines. Microarrays analysis of fungal transcripts (Lahrmann et al., 2013, Lahrmann et al., 2015, Zuccaro et al., 2011) and qPCR experiments during *H. vulgare* and *A. thaliana* colonization, showed that PIIN_01005 is induced during root colonization in both experimental hosts but not in CM (Table S1 and Fig. 2D), complementing the LC-MS/MS analysis. Additionally, the expression of this gene responds to the presence of nucleotides in the medium (Fig S1D). A C-terminal GPI anchor for PIIN_01005 was predicted by the program FragAnchor (Poisson, Chauve et al., 2007). The presence of this protein in the apoplastic fluid of barley indicates that a soluble form might originate from cleavage of the anchor by phosphatidylinositol-specific phospholipase or by proteolytic cleavage as suggested for some animal 5´-nucleotidases (Fini, Talamo et al., 2003, Klemens, Sherman et al., 1990, Vogel, Kowalewski et al., 1992).

Differences in oligomeric nature of E5’NTs were suggested to results in differences in substrate specificity (Knapp, Zebisch et al., 2012). Animal E5’NTs are dimeric, whereas bacterial E5’NTs are monomeric and have a broad substrate specificity being able to hydrolyze ATP, ADP, AMP and other 5’-ribo and 5’-deoxyribonucleotides (Knapp et al., 2012, Thammavongsa, Missiakas et al., 2013). Little is known about the activity of fungal ecto-5´NTs (Russo-Abrahao, Cosentino-Gomes et al., 2011). Since the oligomeric nature of the protein seems to play an important role in substrate specificity, we carried out a prediction of oligomerization based on sequence alignment with the available proteins structures (Fig. 3A, B, C). The crystal structure of the human E5’NT shows that the loop (Cys476–Pro482) is crucial for dimer formation (Knapp et al., 2012). Sequence alignment and structure predictions showed that this loop region is absent in the bacterial 5’NTs and in the *Si*E5’NT (Fig. 3A), suggesting that the fungal protein is monomeric and likely capable of hydrolyzing different nucleotides.

**Figure 3.**
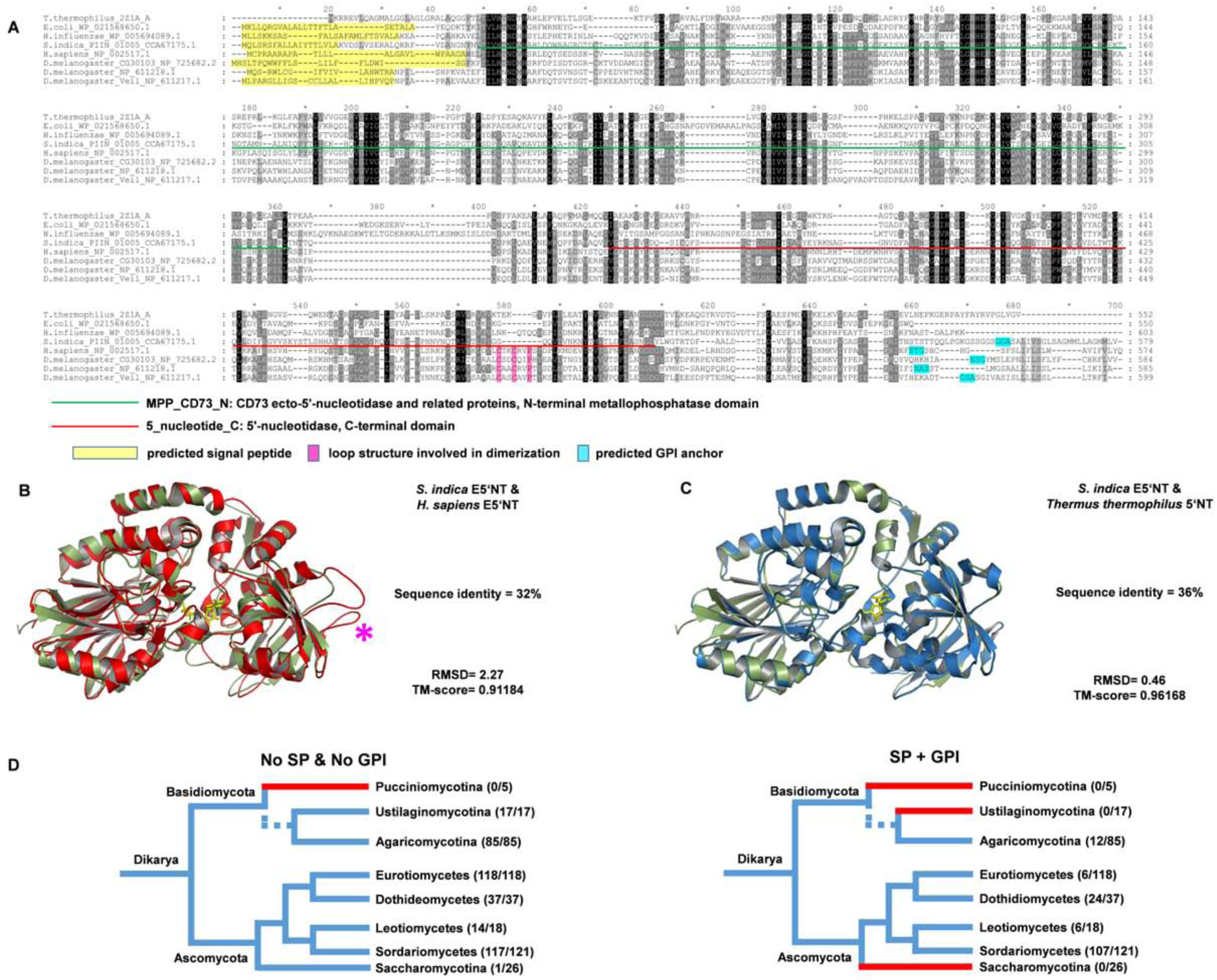
*S. indica* PIIN_01005 encodes a secreted ecto-5’-nucleotidase (*Si*E5’NT). **A** Sequence alignment of *S. indica,* bacterial and animal 5’NTs. *Si*E5’NT and animal E5’NT contain GPI anchors (predicted by FragAnchor; turquoise) and animal E5’NT a loop involved in dimerization (pink). **B** Comparison of the *Si*E5’NT structural homology model (green) with the crystal structure of human E5’NT (PDB id 4H2I, red) with 32% sequence identity. **C** Comparison of the *Si*E5’NT structural homology model (green) with the crystal structure of *Thermus thermophiles* 5’NT (2Z1A, blue) with 36% sequence identity. **\***: position of the loop involved in dimerization of human E5’NT. **D** Distribution of *Si*E5’NT orthologues across higher fungi. End nodes are color-coded based on the presence (blue) or absence (red) of 5’NT genes in a particular fungal taxa. Numbers in parentheses besides the nodes specify the number of species that have 5’NT genes with respect to the total number of genomes analyzed. Left tree: distribution of 5’NT without signal peptide (SP) and GPI anchor. Right tree: E5’NT with SP and GPI anchor. The distribution shows that E5’NT genes are mostly present in Ascomycota such as Sordariomycetes (107/121), followed by Dothideomycetes (24/37) and finally Leotiomycetes (6/18). Few species of Eurotiomycetes (6/118) possess an E5’NT orthologue. In Basidiomycota, E5’NT members are only found in the class of Agaromycotina (12/85).

In order to functionally evaluate the substrate specificity, the full length *S. indica* E5’NT gene was expressed in *A. thaliana* and in the phytopathogenic fungus *Ustilago maydis*. Both organisms have no predicted gene encoding a secreted E5’NT in their genomes. Ecto-5´-nucleotidase activity was determined by the rate of inorganic phosphate released after incubation of 50 µM or 100 µM of either ATP, ADP or AMP with washed culture suspensions of *U. maydis* and with preparations of plasma membrane proteins from *A. thaliana* transgenic lines respectively. Intact cells of the *U. maydis* control strains ^Potef^::mCherry, ^Potef^::E5´NT^woGPI^ and ^Potef^::E5´NT^woSPwoGPI^ were able to hydrolyze ATP and ADP whereas hydrolysis of AMP occurred at very low rate. The *U. maydis* ^Potef^::E5´NT strain expressing the full length *S. indica* ecto-5´-nucleotidase showed significantly higher hydrolysis rates for adenylates, including AMP, compared to the control strains. This indicates that similar to the monomeric bacterial variant, *Si*E5´NT has a broad substrate specificity (Fig. S4) and its activity is extracellular. This was confirmed by the activity measurements of ectopically expressed *Si*E5´NT in *A. thaliana*. Three *A. thaliana* transgenic lines in Col-0 background expressing either ^Pro35S^::E5´NT (303 1L3), ^Pro35S^::SP_E5NT_:mCherry:E5´NT^woSP^ (304 6L8) or ^Pro35S^::mCherry (305 2L1) were selected based on their expression levels for the exogenous gene in the T3 generation (Fig. S5A, B). Ecto-5´-nucleotidase enzyme activity in the membrane fractions demonstrated that expression of the native form of *Si*E5´NT in *A. thaliana* leads to the production of a membrane located enzyme capable of hydrolyzing ATP, ADP and AMP (Fig. 4A). The membrane fraction of the *A. thaliana* line producing *Si*E5´NT-mCherry fusion did not have an increased membrane-bound activity compared to the mCherry control line, suggesting that C-terminal fusion of the *Si*E5´NT to mCherry may reduce its membrane-bound activity. No activity on AMP was detected for *A. thaliana* ^Pro35S^::SP_E5NT_:mCherry:E5´NT^woSP^or^Pro35S^::mCherry lines, implying that no E5´NT-like activity are naturally present under this growth conditions in *A. thaliana*. The *A. thaliana Si*E5´NT plants were otherwise morphologically similar to the control lines in having the same fresh shoot weight (Fig. S5C, D) but they produced less seeds (Fig. S5E).

**Figure 4.**
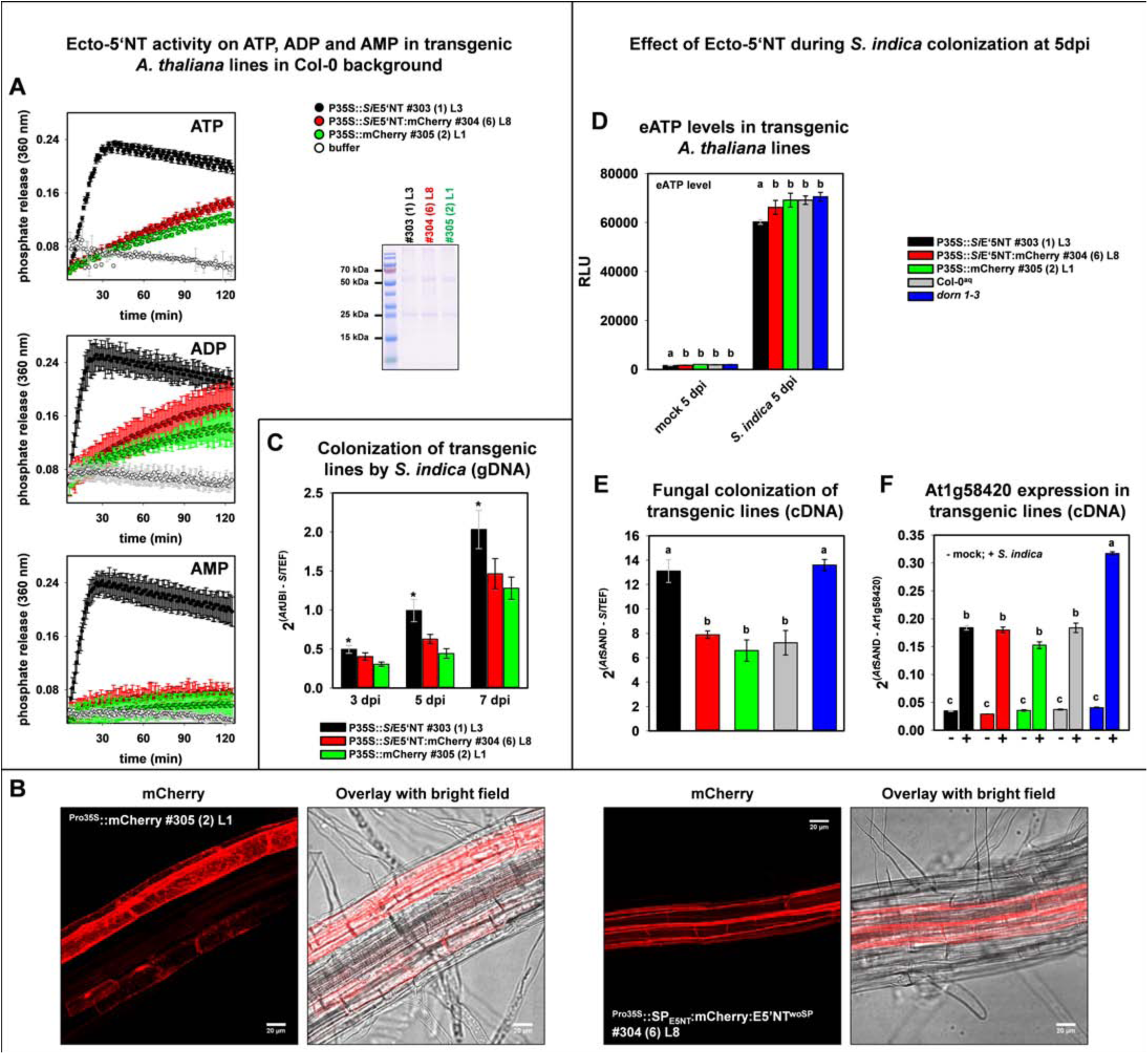
The ecto-5’-nucleotidase activity of *Si*E5’NT in the apoplast of *Arabidopsis* leads to an increase in *S. indica* colonization similar to that of the *dorn1-3* (ATP receptor) mutant line. **A** Ecto-5’-nucleotidase activity measured in membrane protein preparations of *Arabidopsis* plants expressing ^Pro35S^::E5´NT (#303), ^Pro35S^::SP_E5NT_:mCherry:E5´NT^woSP^ (#304) or ^Pro35S^::mCherry (#305). E5’NT activity was measured after incubation with 100 µM of either ATP, ADP or AMP. In the membrane protein preparations from ^Pro35S^::E5´NT (#303) lines phosphate release was specifically increased upon incubation with purines. Error bars represent the standard error of the mean from three technical repetitions. The coomassie stained SDS-PAGE shows the protein pattern of the membrane fractions for the individual transgenic lines. Equal volumes were loaded. The experiment was repeated two times with similar results. **B** Confocal microscopy images of *Arabidopsis* roots expressing either cytosolic mCherry (#305) or ^Pro35S^::SP_E5NT_:mCherry:E5´NT^woSP^ (#304) showing secretion of the E5’NT fusion protein. mCherry images show z-stacks of 14 image planes of 1 µm each. Scale bar=20 µm. **C** The transgenic *Arabidopsis* line ^Pro35S^::E5´NT (#303) expressing untagged full length *Si*E5’NT was better colonized by *S. indica.* Error bars of the qPCR data represent ±SE of the mean from three independent biological replicates. Asterisks indicate significance (Student’s t-test, * p< 0.05). **D** *S. indica* induced eATP release in different *Arabidopsis* transgenic lines. Culture medium was collected from mock treated or *S. indica* inoculated seedlings growing in liquid medium at 5 dpi and released eATP was measured. RLU: relative light units. Error bars represent ±SE of the mean from three biological replicates. Asterisks indicate significance to all other samples within the same treatment group (ANOVA, p<0.05). **E** *S. indica* colonization of transgenic lines at 5 dpi. **F** Expression analysis of the eATP responsive gene At1g58420 measured by qRT-PCR. Error bars represent ±SE of the mean from three independent biological replicates (independent from those shown in Fig. 2C). Letters indicate significant groups (ANOVA, p<0.01, for the line 305 p<0.05).

To confirm secretion of this fungal enzyme in *A. thaliana* transgenic lines, confocal microscopy was carried out with the *Si*E5´NT-mCherry (304 6L8) line. A fluorescence signal in the red channel was visible at the periphery of the plant cells and not in the cytoplasm (Fig. 4B, S5F). Arabidopsis roots were treated with a 0.5 M sorbitol solution to trigger effusion of water from the cytoplasm leading to a volumetric extension of the apoplastic space and plasmolysis. After plasmolysis, the fluorescence signal in the red channel was visible at the membrane and cell wall (Fig. S5G). No auto-fluorescence signals were detected in the UV channel and in the control Col-0 WT line (Fig. S5F, G).

### SiE5’NT enhances fungal colonization and affects eATP levels and defense signaling

To investigate the role of *Si*E5’NT in fungal accommodation, the *A. thaliana* line 303 1L3 constitutively expressing the enzymatically active *Si*E5’NT (Fig. 4A) was used along with the line ^Pro35S^::SP_E5NT_:mCherry:E5´NT^woSP^ 304 6L8 and the control line ^Pro35S^::mCherry 305 2L1 for colonization assays at 3, 5 and 7 dpi (Fig. 4C and S6A, B, C). We observed a significantly enhanced fungal colonization in the *Si*E5’NT plants with respect to *A. thaliana* lines with no enhanced ecto 5’NT enzyme activities (Fig. 4C) at all time points. Complementary, the *Si*E5’NT plants had significant lower level of free eATP than the control lines at 5 dpi were we observed the highest levels of eATP accumulation upon fungal colonization (Fig. 4D). Interestingly, mutation of the DORN1 receptor did not affect eATP concentrations (Fig. 4D), suggesting that perception of eATP is not involved in the regulation of its extracellular levels.

To define the transcriptional response of the *A. thaliana* transgenic *Si*E5’NT line to fungal colonization, qPCR analyses were performed at 5 dpi with AT1g58420, a marker gene reported to be induced by ATP and wounding (Choi et al., 2014). Mutation of the DORN1 receptor was shown to abolish ATP-and to reduce wounding-induced gene expression (Choi et al., 2014). In our experiments AT1g58420 was highly responsive to *S. indica* colonization (Fig. 4F) and (Lahrmann et al., 2013, Lahrmann et al., 2015, Zuccaro et al., 2011). The *dorn1-3* mutant plants displayed increased colonization of *S. indica,* which correlated to the increased expression of the marker gene At1g58420 compared to the colonized control lines (Fig. 4E, F). This implicates that induction of this marker gene by *S. indica* is not solely related to increased ATP levels and its detection by DORN1 in the apoplast. Despite a similar level of fungal colonization (Fig. 4E), the *Si*E5’NT plants had a significantly lower induction of this marker gene compared to the *dorn1-3* mutant plants (Fig. 4F). Expression of two additional marker genes responsive to ATP, CPK28 and RBOHD were also analyzed. These genes showed a similar pattern as observed for the AT1g58420 marker gene with lower induction for the *Si*E5’NT plants compared to the *dorn1-3* mutant plants, but their induction in response to fungal colonization was not as high as for the marker gene AT1g58420 (Fig. S7). These genes are also responsive to different microbe-associated molecular patterns (MAMPs) and microbial treatments, including chitin and flagellin (Fig. S8, and public microarrays, GENEVESTIGATOR, https://genevestigator.com/gv/). To test if *Si*E5’NT would also increase susceptibility to fungal disease, we additionally assessed colonization of Arabidopsis roots by the pathogen *Colletotrichium incanum* which attacks members of the Brassicaceae, Fabaceae and Solanaceae and severely inhibits Arabidopsis growth (Hacquard, Kracher et al., 2016). The *Si*E5’NT plants had a significantly higher colonization by this pathogen, confirming the immunosuppressive function of this fungal enzyme (Fig. S9). Indeed also some pathogenic fungi, including *C. incanum* as well as endophytes like *C. tofieldiae (Hacquard et al., 2016, Hiruma, Gerlach et al., 2016)*, possess a secreted E5’NT which is induced during infection.

Taken together our data demonstrate that eATP accumulates in the apoplast upon *S. indica* colonization at early time points and functions as a danger signal, which is recognized by the DORN1 receptor leading to an induction of defense genes. Extracellular *Si*E5’NT activity leads to significant decreased amount of free eATP in the system and to suppression of defense signaling. Production of bioactive nucleoside intermediates such as adenosine, might also have a role in counteracting ATP-induced signaling as described in animal systems and recently in Arabidopsis (Daumann, Fischer et al., 2015, Matsuoka & Ohkubo, 2004, Zimmermann, Zebisch et al., 2012). In addition, the phosphate released from the complete hydrolysis of eATP could serve as a nutrient. It was shown that *S. indica* transfers phosphate to the host and improve plant growth more effectively at low phosphate compared to high phosphate conditions. Improved growth is observed at late colonization stages but not in the seedlings (Kumar, Yadav et al., 2011, Yadav, Kumar et al., 2010). To evaluate the effect of an additional apoplastic Pi-source on the nutrient exchange between plant and fungus, we employed a computational cell biology approach and simulated the dynamics of a network of proton pumps and proton-coupled transporters during the biotrophic interaction phase. We first used a model where the plant provides sugar to the fungus while the fungus provides phosphate to the plant as described for classical mycorrhizal associations (Schott, Valdebenito et al., 2016). When apoplastic ATP levels and subsequent Pi levels increase, the computational simulations predict that the sugar flux from the plant to the fungus is not affected but the phosphate fluxes change. An external Pi-source provokes an increased Pi-uptake by the plant and a reduced Pi-release or even an uptake of Pi by the fungus (Fig. S10). Thus, the hydrolysis of eATP to adenosine and phosphate could also serve for the Pi-nutrition of the fungus in the early colonization phase without affecting sugar transfer. This implies that the activity of *Si*E5’NT in the apoplast could serve both, nutritional needs and modulation of host immunity.

## Discussion

To establish an integrated and holistic view of the role of the metabolic interplay between plants and their microbes it is of paramount importance to investigate the mechanisms by which plant-associated microbes manipulate plant derived metabolites and how plant metabolism is linked to immunity. Apoplastic communication and its metabolic fluxes have crucial functions in mediating microbial accommodation. During root colonization, plants and microbes secrete a number of proteins to the apoplast that are important for the outcome of the interaction. Plant cells release small amounts of ATP into their extracellular matrix as they grow (Roux & Steinebrunner, 2007). The eATP level can modulate the rate of cell growth in diverse tissues. Beside the physiological role in growth modulation, eATP is released in the extracellular environment in response to biotic stresses modulating defense responses e.g. upon wounding and also when their plasma membranes are stretched during delivery of secretory vesicles (Clark et al., 2014). This implicates a regulatory role for enzymes that can hydrolyze extracellular nucleosides and thus limit their accumulation in the extracellular environment. In plants levels of extracellular nucleotides is controlled by various phosphatases, prominent among which are apyrases EC 3.6.1.5 (nucleoside triphosphate diphosphohydrolases, NTPDases) (Clark et al., 2014). In animals, ecto-nucleotidases play a pivotal role in purinergic signal transmission where the major extracellular purine and pyrimidine compounds known to elicit cell surface receptor-mediated signals are ATP, ADP, UTP, UDP, UDP-glucose, and some additional nucleotide sugars, some dinucleoside polyphosphates, and the nucleoside adenosine (Burnstock, 2007, Burnstock, 2012). The four major groups of ecto-nucleotidases involved in extracellular purinergic signaling include the ecto-nucleoside triphosphate diphosphohydrolases (E-NTPDases), ecto5′-nucleotidase (eN), ecto-nucleotide pyrophosphatase/phosphodiesterases (E-NPPs), and alkaline phosphatases (APs) (Zimmermann et al., 2012). Depending on subtype, ecto-nucleotidases typically hydrolyze nucleoside tri-, di-, and monophosphates and dinucleoside polyphosphates producing nucleoside diphosphates, nucleoside monophosphates, nucleosides, phosphate, and inorganic pyrophosphate (PPi) (Zimmermann et al., 2012). The extracellular membrane-bound ecto-5′-nucleotidase CD73 is anchored to the plasma membrane via a glycosyl phosphatidylinositol (GPI) anchor (Strater, 2006, Zimmermann et al., 2012). This enzyme hydrolysis AMP to adenosine where ADP and ATP are competitive inhibitors, with inhibition constants in the low micro molar range (Knapp et al., 2012). The CD73 together with the ecto-apyrase CD39, that catalyzes the phosphohydrolysis of ATP and ADP to AMP, are part of a cascade to terminate the action of nucleotides as extracellular signaling molecules by decreasing the levels of eATP and eADP and by generating adenosine. In animals, this latter bioactive nucleoside can activate the purinergic ligand-gated ion channel P2X and the G protein–coupled P2Y receptors counteracting ATP-induced signaling (Zimmermann et al., 2012). A purinergic receptor for adenosine or other bioactive nucleoside intermediates is not known in plants and its presence at the host cell surface needs to be proven. Yet it is tempting to speculate that also in plants a similar mechanism is in place where metabolites derived from the hydrolysis of nucleotides modulate the ATP activated immunity once the signaling is not required anymore.

5′-nucleotidases have been found in bacteria, animals and fungi and display significant differences in the range of substrates hydrolyzed and localization. Most fungi have at least one copy of a cytosolic 5´NTs without predicted signal peptide and GPI anchor, whereas secreted ecto-enzyme members are discontinuously distributed within the fungal kingdom (Fig. 3D). E5´NT members are not represented in the available genomes for the Ustilaginomycotina, Pucciniomycotina and Saccharomycotina and only rarely found in the Eurotiomycetes. However, they are widely distributed in two fungal Ascomycota classes, the Sordariomycetes with 107 from the 121 taxa exanimated and the Dothidiomycetes with 24 from 37 taxa. In the Basidiomycota E5´NT members are only present in the Agaromycotina (Fig. 3D). These fungal classes are among the largest groups of fungi with a high level of ecological diversity including many plant and insect pathogens infecting a broad range of hosts.

LC-MS/MS, cytological and transcriptional analyses show induction and secretion of *S. indica* E5´NT during colonization of roots at different symbiotic stages. Functional characterization of the substrate specificity of full-length *Si*E5´NT demonstrated that this fungal enzyme hydrolyzes different nucleotides extracellularly. The apoplastic *Si*E5´NT is sufficient to affect the colonization potential of several fungal species. C-terminal fusion of the *Si*E5´NT to mCherry seems to reduce its membrane-bound activity and its effect on fungal colonization. These data show that during *S. indica* colonization the extracellular *Si*E5´NT activity alone would be sufficient to counteract the action of the bioactive nucleotides ATP and ADP on the DORN1 receptor (Fig. S3). This is supported by the fact that the roots of *A. thaliana* E5´NT expressing line is better colonized by fungi compared to the control lines, accumulates significantly less eATP upon fungal challenge and has lower expression levels for ATP and fungal responsive genes (Fig. 4). In conclusion our work demonstrate that eATP perception plays an important role in plant-fungal interaction in the roots and that *S. indica* has an intrinsic mechanism to counteract bioactive extracellular nucleotides-mediated host defense (Fig. 5). Modelling of P/C exchange between the plant and the fungus in the presence of an apoplastic Pi source derived from eATP hydrolysis suggests that in the early phase of interaction the fungus could reduce phosphate transfer to the plant without affecting the carbohydrate flux from the plant and eventually use this phosphate as a nutrient to support its own growth. Secreted E5´NT homologues are present also in fungal pathogens as *C. incanum* and other endophytes of Arabidopsis such as *C. tofieldiae* where their expression is induced during colonization. Fungal genes encoding enzymes involved in ATP scavenging were also found in the apoplastic fluid of rice leaves infected with the rice-blast pathogens *M. oryzae* and *C. miyabeanus* (Kim, Wu et al., 2014, Kim et al., 2013), suggesting that extracellular purines-based biomolecules play a role also in other plant-fungus interactions. Hydrolysis of plant derived primary metabolites with immune modulating functions in the apoplast could thus represent a general mechanisms by plant-associated fungi to overcome plant defense in addition to serve nutritional needs.

**Figure 5.**
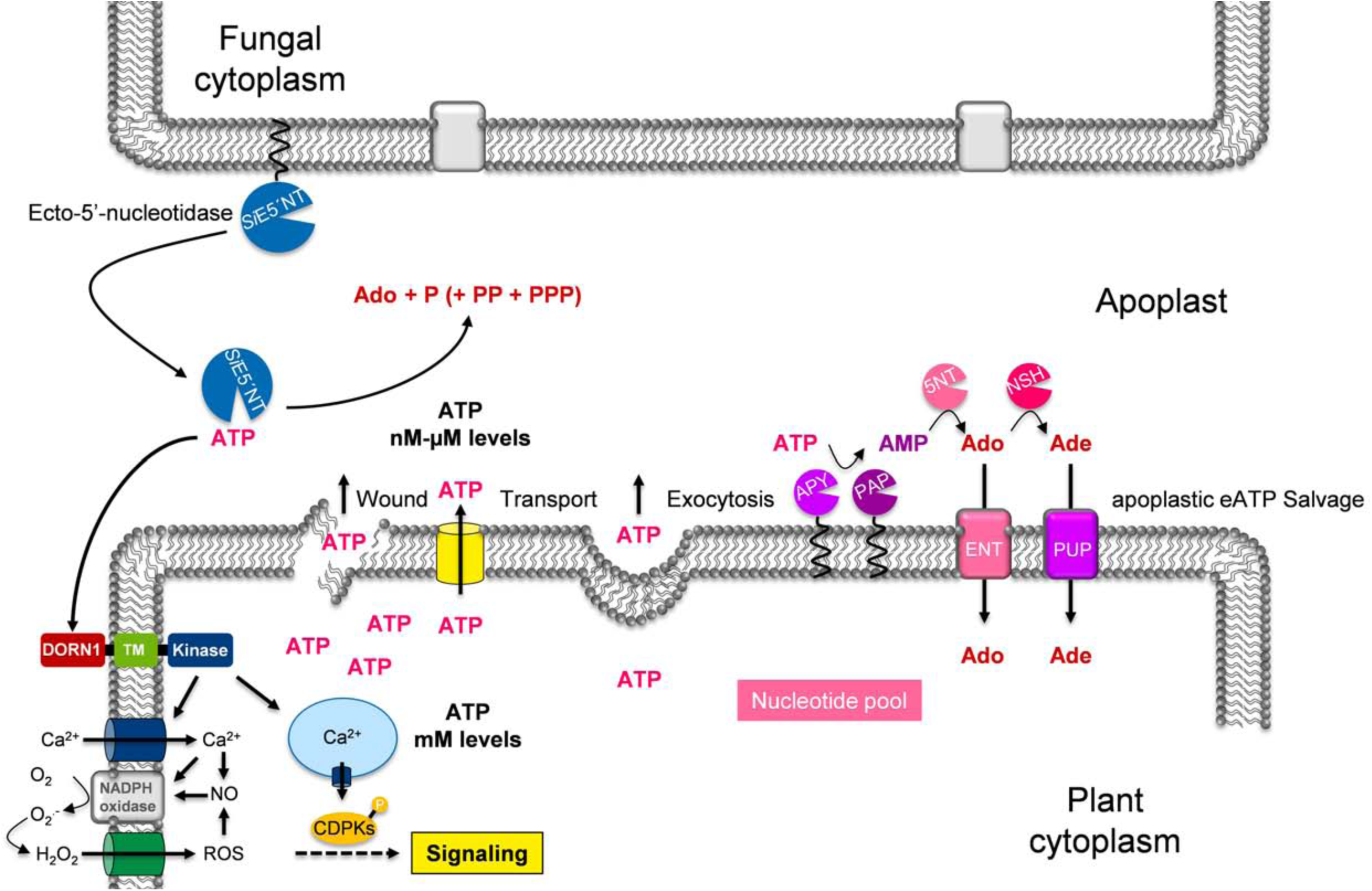
Schematic model showing the potential interference of *Si*E5’NT with apoplastic eATP signaling and salvage pathway in *Arabidopsis*. The schema was modified from (Tanaka et al., 2014). ENT = equilibrative nucleoside transporters, PAP = purple acid phosphatases, Apy = ectoapyrases or nucleoside triphosphate diphosphohydrolases (NTPDases), NSH = nucleoside hydrolases, 5NT = hypothetical 5′-nucleotidases, PUP = hypothetical purine permeases

## Material and methods

### Fungal strains and culturing techniques

*Serendipita indica* (syn. *Piriformospora indica*, DSM11827, Leibniz Institute DSMZ-German Collection of Microorganisms and Cell Cultures, Braunschweig, Germany) and *Serendipita indica* GoGFP strain expressing cytosolic GFP (Hilbert et al., 2012b) were grown in liquid CM supplemented with 2% glucose and incubated at 28°C with constant agitation at 120 rpm, or on solid medium supplemented with 1.5% agar (Basiewicz, Weiss et al., 2012). *Ustilago maydis* SG200 (Kamper, Kahmann et al., 2006) cultures were propagated on potato-dextrose-agar (Difco) or YEPS_Light_ liquid medium at 28°C (Molina & Kahmann, 2007). *Colletotrichum incanum* strain MAFF238704 was kindly provided by Prof. Paul Schulze-Lefert, MPIPZ, Cologne and propagated on solid CM supplemented with 1.5% agar in darkness at 25°C (Hacquard et al., 2016).

### Plant growth conditions and fungal inoculation

Barley seeds (*Hordeum vulgare* cv. Golden Promise) were surface sterilized as described previously (Hilbert et al., 2012b) and germinated for 3 days in the dark on wet filter paper. Germlings were transferred into jars with 1/10 PNM agar (Basiewicz et al., 2012) and inoculated with 3 ml (500,000 spores/ml) *S. indica* chlamydospores in 0.002% Tween20 aqueous solution and cultivated in a growth chamber with a day: night cycle of 16 h : 8 h (light intensity 108 µmol m^−2^s^−1^ and temperature of 22°C: 18°C). Tween water-treated germlings were used as control. Three independent biological replicates were used. For each biological replicate, one jar containing 4 to 5 barley seedlings were pooled at each time point.

*Arabidopsis* seeds were surface sterilized with 70% ethanol for 10?min followed by 100% ethanol for 7 min and then air dried. The sterilized seeds were transferred to half-strength Murashige and Skoog (½MS) medium containing 1% sucrose and 0.4% Gelrite (Duchefa), and incubated for 3 days in the dark at 4 °C and subsequently grown for 7 days with a day: night cycle of 8h: 16h (light intensity, 47 μmolm^−2^s^−1^) at 24°C. Three independent biological replicates were used for all *Arabidopsis* experiments unless otherwise stated in the legend. For each biological replicate, 3 plates containing 20 plants each, were pooled. For colonization studies, (500,000 spores/ml) of *S. indica* chlamydospores in 0.002% Tween20 aqueous solution were applied to the roots of each plant of 7-day old seedlings. Tween water-treated seedlings were used as control.

For colonization analysis using *C. incanum Arabidopsis* seedlings were grown for 10 days under a day: night cycle of 8 h: 16 h regime (light intensity 111 μmolm^−2^s^−1^ and temperature of 22 °C: 18 °C). Roots were drop-inoculated with 200,000 spores/ml of *C. incanum* suspended in sterile distilled water. Water-treated seedlings served as control. Three independent biological replicates were used each representing a pool of roots from 20 plants.

### Identification of *S. indica* proteins in apoplastic fluid and culture filtrate

Barley seeds (*Hordeum vulgare* L. cv Golden Promise) were surface sterilized for 1 min under light shaking in 70% ethanol followed by 1 hour in 12% sodium hypochlorite. Subsequently, seeds were washed repeatedly for 1 hour with sterile distilled water. After germination for three days at 22°C in complete darkness on wet filter paper, germinated seeds were transferred into jars with 1/10 PNM (Basiewicz et al., 2012) under sterile conditions and inoculated with 3 ml of chlamydospores (concentration 500,000 spores/ml) of *S. indica* GoGFP strain (Hilbert, Voll et al., 2012a) and cultivated in a growth chamber with a day: night cycle of 16 h: 8 h (light intensity, 108 µmol m2/s) and temperature of 22°C: 18°C. For APF extraction three time points reflecting different interaction stages have been chosen: biotrophic phase (5 dpi) as well as cell death associated phase (10 and 14 dpi). After respective growth time, roots from 300 seedlings for each treatment and time point were thoroughly washed and cut into pieces of 2 cm length (2nd to 4th cm of the root). Samples were vacuum infiltrated (Vacuumbrand, CVC 3000, VWR) with deionized water 5×15 min at 250 mbar with 1 min atmospheric pressure break. Bundled infiltrated roots were centrifuged in 5 ml syringe barrels at 2000 rpm in a swing bucket rotor (800 g) for 15 min at 4°C as described in (Wawra et al., 2016). Pooled APF samples were stored at −20°C. In order to exclude cytoplasmic contamination apoplastic fluid samples were tested in immunoblots with GFP antibody. For western blotting, proteins from unstained gels were transferred onto nitrocellulose membranes using a semi dry blotting system from Bio-Rad (Hercules, USA). Anti-GFP antibody was obtained from Cell Signalling Technology (Danver, USA) and used at a 1:2000 dilution. An anti-mouse antibody purchased from Sigma (Munich, Germany) was used as secondary antibody at a dilution of 1:2000. For higher peptide mass accuracy part of the APF samples were deglycosylated with Protein Deglycosylation Mix (P6039S, New England BioLabs, Ipswich, USA) under denaturing conditions. In agreement with (Yu, Tang et al., 1999), at the centrifugation force used the samples were free from obvious symplastic contamination, and were thus regarded as being of apoplastic origin.

Secreted proteins from culture filtrate were prepared as follows: *S. indica* cultures grown for 7 days in liquid CM were passed through a miracloth filter (Merck Millipore, Darmstadt), a folding filter type 600P (Carl Roth, Karlsruhe, Germany) and finally a 0.45 µm syringe filter (Millex-GP; Merck Millipore, Darmstadt, Germany) to remove small mycelium fragments and spores. Proteins were precipitated by addition of 10% (v/v, final concentration) trichloroacetic acid and incubation at −20°C overnight and subsequent centrifugation at 40,000 g. For SDS-PAGE, pelleted secreted proteins were resolved in 50 µL of 1X SDS sample buffer and 15 µL were used. For mass spectrometric analysis, pelleted secreted proteins were solved in 50 µL TBS (50 mM Tris pH 7.5, 150 mM NaCl). Half of the samples were deglycosylated with the Protein Deglycosylation Mix (New England Biolabs, Ipswich, USA) under denaturing conditions according to manufacturer’s protocol. Proteins were then analyzed via liquid chomatopgraphy electron spray ionization tandem mass spectrometry (LC-ESI-MS/MS; LTQ Orbitrap Discovery; Thermo Fisher Scientific, Waltham, USA) after tryptic digest (FASP^TM^ Protein Digestion Kit; Expedeon, Swavesey, UK) in the facilities of CEACAD/CMMC Proteomics Facility in Cologne, Germany. Peptide masses were compared to *S. indica in silico* trypsin digested proteome (Genbank, NCBI) with an inclusion of GFP, with a barley dataset downloaded from the IPK Gatersleben homepage (http://webblast.ipk-gatersleben.de/barley_ibsc/downloads/) and with *Rhizobium radiobacter* F4 (syn. *Agrobacterium tumefaciens*, syn. *Agrobacterium fabrum*) dataset downloaded from Genbank, NCBI (WGS project JZLL01000000, https://www.uniprot.org/proteomes/UP000033487). The exact instrument settings can be found in the supplementary Table 6 containing the processed data. Mass spectrometric raw data were processed with Proteome Discoverer (v1.4, Thermo Scientific) and SEQUEST. Briefly, MS2 spectra were searched against the respective databases, including a list of common contaminants. The minimal peptide length was set to 7 amino acids and carbamidomethylation at cysteine residues was considered as a fixed modification. Oxidation (M) and Acetyl (Protein N-term) were included as variable modifications. The match-between runs option was enabled. Plant derived proteins (data not shown) and fungal derived proteins matched to peptides were analyzed for the presence of signal peptides and exclusiveness for the three specific time points of APF extraction.

### Construction of plasmids

Full length *E5’NT* (PIIN_01005) was amplified from cDNA of barley roots inoculated with *S. indica* at 5 dpi by respective oligonucleotides (Table S5) and cloned into the TOPO^®^ cloning plasmid (Thermo Fisher Scientific, Waltham, USA) resulting in TOPO-E5’NT. For the expression of various *E5’NT* constructs under the control of the synthetic otef promoter (Wang, Berndt et al., 2011), *E5’NT* full length, *E5’NTwoGPI and E5’NTwoSPwoGPI* were amplified from the plasmid TOPO-E5’NT using Phusion DNA polymerase (NEB) by respective oligonucleotides (Table S5), and cloned into the *Bam*HI-*Not*I digested plasmid ^Potef::^Yup1(RFP)_2_ (Lenz, Schuchardt et al., 2006) using Gibson assembly (Gibson, Smith et al., 2010), generating the plasmids ^Potef::^E5’NT, ^Potef::^E5’NT^woGPI^ and ^Potef::^E5’NT^woSPwoGPI^, respectively.

For the construction of SP_E5’NT_:mCherry:E5’NT^woSP^, *Si*E5’NT without signal peptide (*E5’NT*^*woSP*^) was amplified from the plasmid TOPO-E5’NT using Phusion DNA polymerase (NEB) by respective oligonucleotides (Table S5), and cloned into the *Bam*HI digested plasmid ^Pro*Um*Pit^2^::^SP_Dld1_:mCherry:Dld1^woSP^ (Nostadt *et al*., unpublished data), generating the plasmid ^Pro*Um*Pit^2^::^SP_Dld1_:mCherry:E5’NT^woSP^. Subsequently, ^Pro*Um*Pit^2^::^SP_Dld1_:mCherry:E5’NT^woSP^ was digested with *Sac*II and *Nco*I and ligated with a DNA fragment encoding the E5’NT signal peptide (SP_E5NT_) digested with the same restriction enzymes, replacing the Dld1 signal peptide (SP_Dld1_) and generating the construct ^Pro*Um*Pit^2^::^SP_E5’NT_:mCherry:E5’NT^woSP^. For expression in *A. thaliana* under the control of CaMV 35S promoter (Pro35S), *E5’NT* full length, SP_E5’NT_:mCherry:E5’NT^woSP^ and mCherry were amplified from the plasmids TOPO-E5’NT and ^Pro*Um*Pit^2^::^SP_E5’NT_:mCherry:E5’NT^woSP^, respectively, using Phusion DNA polymerase (NEB) by respective oligonucleotides (Table S5), and cloned into the *Bam*HI digested plasmid pCXSN (Chen, Songkumarn et al., 2009), using Gibson assembly, generating the plasmids ^Pro35S^::E5’NT, ^Pro35S^::SP_E5’NT_:mCherry:E5’NT^woSP^ and ^Pro35S^::mCherry, respectively.

### *Ustilago maydis* transformation

The *U. maydis* strains were transformed by integration of the p123 derived plasmids into the ip locus in the haploid solopathogenic strain SG200 as previously described (Loubradou, Brachmann et al., 2001).

### *Ustilago maydis* cell surface 5’-nucleotidase activity assay

For the cell surface 5’-nucleotidase activity assay, ^Potef^::E5’NT, ^Potef^::E5’NT^woGPI^^Potef^::E5’NT^woSPwoGPI^ along with the control ^Potef^::mCherry *U. maydis* strains were grown overnight in YEPS_Light_. Then the cultures were diluted in PD broth and grown until OD_600_ reached 0.8. The cells were harvested through centrifugation and washed two times with 0.9% NaCl and finally re-suspended in equal volume of sterile double deionized water. Then 100 µl of cell suspension from each *U. maydis* strains was incubated with 50 µM of ATP, ADP or AMP for 30 min, and the hydrolysis of these nucleotides was measured by quantifying the amount of phosphate released using EnzChek^®^ Phosphate Assay Kit (Thermo Fisher Scientific).

### *A. thaliana* transformation

The plasmids ^Pro35S^:: E5´ NT(#303), ^Pro35S^:: SP _E5NT_:m Cherry: E5´NT^woSP^ (#304) and ^Pro35S^::mCherry (#305) were transformed into the *Agrobacterium* strain GV3101 by electroporation using Gene Pulser Xcell Electroporation system (Bio-Rad Laboratories, Hercules, CA, USA) following the manufacture indications. The cells were plated on LB medium containing 25 μg/ml of rifampicin and 50 μg/ml of kanamycin. *Agrobacterium*-mediated transformation of *A. thaliana* was carried out by means of the floral dip method (Clough & Bent, 1998, Zhang, Henriques et al., 2006). Stratified seeds from T0 plants were grown in ½ MS containing 30mg/L hygromycin under short day condition. Transformants were selected by their hygromycin resistance. Putative transformants were transferred to soil and two weeks later tested by PCR using a gene-specific primer (Table S5). The T_3_ transformants from ^35Spro^::E5´NT (#303) and ^35Spro^:: SP_E5NT_: mCherry: E5´NT^woSP^ (#304) were confirmed again by both hygromycin resistance and PCR using a gene-specific primer. In addition, the elevated expression level of *E5’NT* in ^35Spro^::E5´NT (#303), ^35Spro^::SP_E5NT_:mCherry:E5´NT^woSP^ (#304) as well as the *mCherry* in ^35Spro^::mCherry (#305) were analyzed by quantitative real-time PCR. The transgenic lines which had the highest transcript levels were selected for further studies.

### ATP assay

To test whether colonization of *S. indica* in *Arabidopsis* and barley triggers the release of eATP, seven days old *Arabidopsis* seedlings and three days old barley seedlings were treated either with a *S. indica* spore suspension (5 x 10^5^ spores / ml) or with 0.01 % tween water for 2.5 hours. The treated seedlings were transferred to 60 mm petri plates (3 plates per treatment). Each plate contained 3 ml of liquid ½ MS (ammonium free) medium and 15 treated seedlings. Subsequently, seedlings were grown for 3, 5, 7 and 10 days post inoculation (dpi) with constant light at 22°C. Three individual samples of 50 µL media were collected from each plate, flash frozen in liquid nitrogen and stored in −80°C until eATP measurements were carried out. The measurement of eATP in the growth media was performed using the ENLITEN ATP Assay System from Promega. Reactions were carried out in 96-well plates and luminescence was detected using a microplate reader (TECAN Safire).

### Membrane Protein Purification

Membrane bound proteins were extracted from 1 g shoot material from the respective *A. thaliana* lines (25 days old) grown in soil. The material was ground in liquid nitrogen and re-suspended in 5 ml 0.05 M Tris/HCl buffer containing 1mM MgCl_2_ (pH 7.5). After centrifugation for 20 min at 12,000 g the pellet was resuspended in 5 ml Tris buffer containing 1% IGEPAL CA-630 (Sigma), gently stirred at 4 °C for 30 min, subsequently centrifuged at 5,000 g for 10 min. The supernatant was cleared by ultracentrifugation at 100,000 g for 45 min. The resulting supernatant was used for subsequent enzyme activity assays.

### Measurement of E5’NT and Apyrase Activity Assay in *A. thaliana* lines

The activity of purified, membrane bound apyrases and the overexpressed *Si*E5’NT was monitored using the EnzChek^®^ Phosphate Assay Kit (Thermo Fisher Scientific) according to manufacturer’s instruction. 15 µl of purified membrane proteins were used for the enzyme kinetics assay. 100 µM ATP, ADP or AMP (in 10 mM Tris/MES buffer containing 2 mM MgCl_2_ and 30 mM KCl; pH 6.5) were used as substrates. The background absorbance at 360 nm was monitored for 5 minutes before the addition of the substrates. Subsequently, kinetics were measured for 2 h.

### Real-Time PCR analysis

DNA isolation was performed from ∼200 mg of ground material. The powder was incubated for 5 min at RT under slight rotation with ∼ 500 µl CTAB buffer (2 % Hexadecyl trimethyl-ammonium bromide, 100 mM Tris/HCl, 20 mM EDTA, 1.4 M NaCl and 1 % polyvinyl pyrrolidone vinylpyrrolidine homopolymer Mw 40,000, pH 5.0). Subsequently, 250 µl chloroform : isoamyl-alcohol (24:1) was added followed by an additional incubation for 5 min at RT and centrifugation. The water phase was collected and polysaccharides were precipitated using ethanol before final DNA precipitation using 1 volume of isopropanol. Total RNA was isolated from 200 mg of ground fungal material or colonized and non-colonized plant root material using the TRIzol^®^ reagent (Invitrogen, USA) as described previously (Hilbert et al., 2012a). Contaminating genomic DNA was removed by DNaseI treatments (Thermo Fisher Scientific, USA) according to manufacturer’s protocol. First-strand cDNA was synthesized with 1 µg of total RNA primed with Oligo-dT and random hexamer using First strand cDNA synthesis Kit (Thermo Fisher Scientific, USA). The resulting cDNA preparations were diluted to a final concentration of 2.5 ng/µl and 4 µl of each cDNA sample was used for qRT-PCR. *S. indica* Translation Elongation Factor1 (*SiTEF*) and *A. thaliana SAND* or *UBI* gene primers were used for qPCR amplifications. The reactions were performed in 96-well reaction plate (BioRad, USA) on a Real-Time PCR System (BioRad, USA). Each well contained 7.5 µl of 2X SYBR^®^ Green QPCR Master Mix (Promega, USA), 4 µl of cDNA or 10 ng of DNA, and 0.7 µl of each primer from a 10 µM stock, in a final volume of 15 µl. Transcripts of each gene were ampli?ed using the primers described in Supplementary Table S5. The threshold cycles (C_T_) of each gene were averaged from three replicate reactions and the relative expression or plant to fungus ratio from gDNA was calculated using the 2^−ΔCT^ method (Livak & Schmittgen, 2001).

### E5’NT expression levels in axenic culture in response to nucleotides

In order to test whether SiE5’NT expression is induced in the presence of nucleotides, WT *S. indica* spores (500,000 spores/ml) were used to inoculate liquid CM medium and grown for 5 days at 28°C with 110 rpm shaking. Subsequently, the culture was filtrated through miracloth, the mycelium was crushed and regenerated in 100 ml fresh CM medium for 2 days. After regeneration, 5 ml of fungal suspension was diluted with 45 ml of CM medium supplemented with 100 µM of the respective nucleotides. At the indicated time points the mycelium was harvested by filtration, washed with water and flash frozen in liquid nitrogen before processing according to the Real-Time PCR analysis protocol listed above. 3-6 independent biological replicates for each time point/nucleotide combination were performed.

### Calcium Influx Quantification

All calcium influx experiments were conducted using plants that ectopically express Aequorin as Ca^2^+ reporter (Choi et al. 2014). 7 days old seedlings grown on ½ MS plates containing 1% sucrose were transferred to 96-well plates containing 50 µl 10 µM Coelenterazine (Promega) in reconstitution buffer (2 mM MES/KOH containing 10 mM CaCl_2_; pH 5.7). After 16 h the luminescence background was measured for 5 min followed by the addition of different adenylates (final concentration 100 µM ATP, ADP, AMP or adenosine). Luminescence was monitored for 30 min.

### Confocal microscopy

Confocal images were recorded using a TCS-SP8 confocal microscope (Leica, Bensheim, Germany). mCherry fluorescence was imaged with an excitation of 561 nm and a detection bandwidth set to 580–630 nm. DAPI and auto fluorescence of the samples were assessed by laser light excitation at 405 nm with a detection bandwidth set to 415-465 nm.

### Homology modeling of *Si*E5’NT

In order to carry out the homology modeling of SiE5’NT, best template was selected through PSI BLAST against the PDB database (http://www.rcsb.org/pdb/home/home.do). The best hit with high sequence identity and atomic resolution < 1.8 Angstroms, was selected as template. The three dimensional structure of SiE5’NT was generated using a restrained-based approach in MODELLER9v11 (Webb & Sali, 2017). Initial refinement of the 3D model generated was carried out with the help of loop refinement protocol of MODELLER. The assessment of the final structural model was carried out with PROCHECK (Laskowski, Macarthur et al., 1993), ProSA (Wiederstein & Sippl, 2007) and QMEAN (Benkert, Kunzli et al., 2009) analyses. The structure was visualized using PyMOL (http://www.pymol.org/).

### Computational Cell Biology

The modelling approach was essentially performed as described previously (1). The minimal model of (Schott et al., 2016) was enlarged by adding the release of ATP from the plant into the apoplast and its subsequent cleavage into phosphates and adenosine by E5’NT enzymes. The mathematical description of these processes results in redundant parameters, which were combined. Therefore, ATP-release and cleavage were considered as one process in this study, which was modelled as a constant Pi-production rate: d[Pi]_apo_ = const×dt. Following the mathematical description of all transporters, an *in silico* cellular system was programmed and computational cell biology (dry-lab) experiments were performed using the VCell Modeling and Analysis platform developed by the National Resource for Cell Analysis and Modeling, University of Connecticut Health Center (Loew & Schaff, 2001).

## Acknowledgment

We would like to thank Prof. Gary Stacey for providing the *A. thaliana dorn1-3* mutant and Col-0^aq^ lines. Heidi Widmer is acknowledged for her excellent support in the lab. Ganga Jeena is acknowledged for her support with bioinformatics analysis. ID was supported by the Fondecyt grant No. 1150054 of the Comisión Nacional Científica y Tecnológica of Chile. AZ acknowledge support from the Cluster of Excellence on Plant Science (CEPLAS, EXC 1028), DFG ZU 263/2-1, DFG ZU 263/3-1 and institutional funds of the Max Planck Society.

## Conflict of Interests

The authors declare that they have no conflict of interest.

## Author contributions

AZ and SN conceived the project, designed the experiments and wrote the paper with contributions from SW and GL. SN, SW, XQ, RN, FG, FS, GL performed experiments and analyzed the data. ID produced the P/C model.

## Supplemental figure legends

**Supplementary figure 1. A** Schematic workflow describing the barley inoculation and apoplastic fluid (APF) collection. To ensure that no apoplastic leakage of the *S. indica* GoGFP strain expressing cytosolic GFP used for the inoculation of the barley seedling occurred during APF collection the samples dedicated for mass spectrometric analysis were examined by an anti-GFP western blot. **B** *S. indica* putative apoplastic proteins involved in metabolism shown within their respective cellular pathways. Asterisks indicate orthologues identified in the APF of rice leaves infected with the rice-blast pathogen *Magnaporthe oryzae* (Kim et al., 2013). The metabolic pathways were modified from (Pechanova, Hsu et al., 2010). **C** Peptides identified by LC-MS/MS for E5’NT found in the APF of barley. BLAST searches against barley databank retrieved no perfect matches, indicating that these peptides originated from *S. indica* E5’NT. **D** Expression of *Si*E5’NT in *S. indica* grown in liquid CM supplemented with 100 µM ATP, ADP, AMP or ATP-imido (adenosine 5′-(β,γ-imido)triphosphate, SIGMA) a non-hydrolysable form of ATP. Mock 0 represents the time point 0 just before addition of the nucleotides and it is set to 1. An induction of expression was detected at 12 hour post treatment for AMP and ATP-imido. Significances obtained by one way ANOVA testing of each individual time point are indicated.

**Supplementary figure 2. Gene Ontology Enrichment Analyses of apoplastic and culture filtrate proteins.**

**Supplementary figure 3. Characterization of the *dorn1-3* mutant line.** ATP and ADP but not AMP nor adenosine induce intracellular calcium release in *Arabidopsis thaliana* expressing aequorin (Col-0^aq^). The ATP receptor deletion mutant *dorn1-3* does not respond to ATP and ADP. After 5 min of background measurements 50 µl of 100 µM adenylates were added, subsequently luminescence was monitored for 30 min. **A**, ATP; **B**, ADP; **C**, AMP; **D**, adenosine. Data points represent means of six independent biological replicates ±SE.

**Supplementary figure 4. Characterization of *Si*E5’NT. A** Cell surface 5’-nucleotidase activity assay of *Ustilago maydis* strains expressing different forms of *Si*E5’NT. To study the ecto-5’-nucleotidase activity of *Si*E5’NT and its substrate preferences, the biotrophic plant pathogen smut fungus *U. maydis* was chosen since an E5’NT orthologue predicted to be secreted is absent in its genome. Ecto-5’-nucleotidase activity of *U. maydis* cells expressing different *Si*E5’NT versions under the control of the constitutive synthetic otef promoter (Wang et al., 2011) were analyzed. ^Potef^::mCherry encodes for cytosolic mCherry, ^Potef^::E5’NT encodes for the full length *S. indica* enzyme, the ^Potef^::E5’NTwoGPI construct lacks the predicted GPI anchor sequence and ^Potef^::E5’NTwoSPwoGPI is missing both the predicted signal peptide and the GPI anchor. Cell surface activity of *Si*E5’NT in *U. maydis* washed cell suspensions were analyzed by measuring the amount of phosphate released after 30 min of incubation with 50 µM of either ATP, ADP or AMP. The amount of phosphate is significantly increased in cells transformed with ^Potef^::E5’NT for all substrates tested. Error bars represent standard deviation of the mean from three biological replicates. Asterisks indicate significance at p< 0.05 (*), 0.01 (**) analyzed by Student’s t-test to the mCherry control strains.

**Supplementary figure 5. Characterization of *A. thaliana* T3 and T4 transgenic lines in Col-0 background. A** *Si*E5’NT and **B** mCherry expression analysis of different *A. thaliana* T3 transgenic lines by qPCR. Error bars represent standard deviation of the mean from three technical replicates of one biological experiment. **C** Shoot weight of *A. thaliana* transgenic T3 lines. Error bars represent standard deviation of the mean from two biological replicates. Arrows indicate the lines that were selected for further analyses. Black indicates E5’NT expressing lines; Red indicates E5’NT:mCherry expressing lines; Green indicates mCherry expressing lines. **D** Images of seedlings grown on ½ MS with Hygromycin B (15 µg/mL) from the T3 transgenic lines used for detailed analyses. **E** Seed production in the *A. thaliana* transgenic T4 lines used for detailed analyses in Figure S9. Plants were propagated in parallel in greenhouse in soil. Seeds from each plants were collected separately and weighted. Twenty to forty plants for each line were used for one way ANOVA statistical analysis. The activity of *Si*E5’NT might affects seed production in the 303 1L3 line by manipulation of eATP levels during propagation. It was reported that optimum concentration of eATP is essential for initiation of pollen germination in *A. thaliana* and in Gymnosperms (Steinebrunner, Wu et al., 2003, Zhou, Fan et al., 2015). **F** Confocal laser scanning microscopy of roots from the lines 304 6L8 expressing E5’NT:mCherry and Col-0 WT under the same settings. **G** Plasmolysis using 0.5 M sorbitol. Plant nuclei were stained with DAPI and visualized with the UV channel. No auto-fluorescence in the UV channel was observed at the laser intensity used. No fluorescence signal was observed in the WT root in the mCherry (red) channel. After plasmolysis a fluorescence signal was detected at the membrane (white arrows) and at the cell wall (yellow arrows), suggesting that E5’NT:mCherry fusion protein is secreted. N indicates a nucleus.

**Supplementary figure 6. A** *S. indica* colonization levels of different independent transgenic T2 and T3 lines. **B** mCherry transcript levels. **C** *Si*E5’NT transcript levels of the respective plants. Samples were analyzed by qPCR. Error bars show the standard error of the mean of three independent biological replicates. Asterisks indicate significance at p< 0.05 (*) analyzed by Student’s t-test to the mCherry control line (#305).

**Supplementary figure 7. *At*RBOHD and *At*CPK28 expression levels upon *S. indica* colonization (addition to Figure 4E/F).** Expression levels of the eATP and wounding responsive genes *At*RBOHD and *At*CPK28 (Choi et al., 2014) in transgenic lines colonized by *S. indica* at 5 dpi. Data were normalized by setting the values from the mock treated samples to 1. Error bars represent ±SE of the mean from three independent biological replicates.

**Supplementary figure 8. Expression patterns of At1g58420, *At*WRKY40, *At*RBOHD and *At*CPK28 under different perturbations, available in the GENEVESTIGATOR database.** The filter for the generation of the dataset was set to a cutoff of fold change >3. These genes are eATP responsive only at low level (below cutoff) and strongly responsive to MAMPs and biotic stresses.

**Supplementary figure 9. Colonization assay of transgenic lines by the phytopathogenic fungus *Colletotrichium incanum*. A** Characterization of the transgenic lines after propagation in the T4 generation. Plants were propagated in parallel in greenhouse in soil. Lines 303 2L3, 304 6L8 and 305 2L1 were chosen for colonization analysis with *C. incanum* based on the expression strength for the respective heterologous gene. *Si*E5’NT transcript levels were measured by qPCR. **B** Colonization of transgenic lines by *C. incanum* at 5 dpi. Error bars show the standard error of the mean of three independent biological replicates. Asterisks indicate significance at p< 0.05 (*) and p< 0.01 (**) analyzed by one way ANOVA.

**Supplementary Figure 10. Simulation of the effect of ATP release and its cleavage in the apoplast on the nutrient exchange between plant and fungus.** The nutrient exchange of phosphates (P) and sugars (C) between plant and fungus in mycorrhizal interactions has been successfully described by Schott et al. (1) in a minimal model combining proton-coupled phosphate (H/P) and sugar (H/C) transporters and proton pump-driven background conductances. This model was enlarged by adding the release of ATP from the plant into the apoplast and its subsequent cleavage into phosphates (Pi) and adenosine (A) by E5’NT enzymes. The Pi and C exchange between the plant and the fungus was simulated for different values of ATP-release and Pi-production. The ATP release-and Pi production-rate was normalized that at the relative value of 1 the fungus does not export Pi. **A** Relative fluxes of Pi and C across the fungal plasma membrane. Without additional Pi-source in the apoplast (value 0 at the x-axis, black dots) there is a constant flux of phosphate via the H/P transporter from the fungus to the apoplast and a constant flux of sugar from the apoplast to the fungus. For better comparison these fluxes were normalized to 1 and-1, respectively, and all other fluxes were calculated relative to these control values. With increasing ATP-release and Pi-production the Pi-efflux gets smaller and is zero at a relative Pi-production rate of 1 (grey triangles). At higher ATP-release and Pi-production rates the fungus imports Pi, i.e. the transport direction of phosphate has been inverted in comparison to the control condition (white square). The C-flux via the H/C transporter is not affected by the additional Pi-source (horizontal grey line). **B** Relative fluxes of Pi and C across the plant plasma membrane. Without additional Pi-source in the apoplast (value 0 at the x-axis, black dots) there is a constant flux of sugar via the H/C transporter from the plant to the apoplast (and thereafter to the fungus) and a constant flux of phosphate (coming from the fungus) via the H/P transporter from the apoplast to the plant. With increasing ATP-release and Pi-production the Pi-influx via the H/P transporter increases while the C-flux via the H/C transporter is unaffected by the additional Pi-source (horizontal grey line). **C** Phosphate fluxes across the plant plasma membrane. Besides the Pi-uptake via the H/P transporter (B, black line, for clarity not shown in C), the plant loses Pi due to the ATP-release (light grey line). The difference between Pi-uptake and Pi-loss is the net Pi-balance of the plant (dark grey line). Without ATP-release (value 0 at the x-axis, black dots) the plant gains Pi. At a relative ATP-release and Pi-production of 1 (grey triangles) the plant release as much phosphate as it takes up, while at higher ATP-release values, the plant loses Pi. **D** Schematic representation of the fluxes for three scenarios: (i) no ATP-release and Pi-production (left, black dots in A-C), (ii) moderate ATP-release and Pi-production (middle, grey triangles in A-C), and (iii) high ATP-release and Pi-production (right, white squares in A-C). If there is no ATP-release (left), there is a constant flux of Pi from the fungus via the apoplast to the plant and a constant flux of sugars in the inverse direction. The energy for these fluxes is provided by the phosphate gradient between fungus and plant and the sugar gradient between plant and fungus. At a moderate ATP-release and Pi-production rate (middle, value 1 on the x-axis, grey triangles in A-C), there is no Pi-flux from the fungus to the apoplast anymore. The unchanged uptake of sugars is now energized by the proton pump of the fungus. In this condition, the plant retrieves the cleaved Pi originating from the ATP-release via the H/P cotransporter, a transport which is energized by both, the sugar gradient and the proton pump. At a high ATP-release and Pi-production rate (right, value 2 on the x-axis, white squares in A-C), there is a larger Pi-uptake by the plant and a Pi-uptake by the fungus. Both processes are energized by larger activities of the proton pumps.

